# Open Fourier Ptychographic Microscopy (OpenFPM)

**DOI:** 10.64898/2026.03.18.711080

**Authors:** Lewis D. Walker, Laura Copeland, Liam M. Rooney, Christopher Bendkowski, Michael Shaw, Gail McConnell

## Abstract

Fourier ptychographic microscopy (FPM) uses sequential multi-angle illumination and iterative phase retrieval to recover a high-resolution complex image from a series of low-resolution brightfield and darkfield images. We present OpenFPM, an open-source FPM platform in which conventional and optomechanical hardware is replaced with compact, low-cost 3D printed components. Illumination, sample and objective positioning, and camera triggering are controlled using a Python-based interface on a Raspberry Pi microcomputer. With a 10 × /0.25 NA objective lens and 636 nm illumination, OpenFPM experimentally achieves amplitude and phase reconstructions with an effective synthetic NA of 0.90 over a 1 mm field-of-view. This platform gives researchers accessible and affordable hardware for developing and testing LED-array microscopy techniques for a range of biomedical imaging applications.

---

Three-dimensional (3D) printing has become an important and accessible tool for low-cost, rapid in-house prototyping in optical and biomedical research [1,2]. Advances in high-precision printing over large build volumes have enabled the development of 3D-printed optical components and opensource microscope platforms that are increasingly used in research applications [3,4]. However, conventional microscopes are fundamentally constrained by the numerical aperture of the objective lens (*NA*_*obj*_). A high *NA*_*obj*_ enables the capture of high spatial frequency information for high-resolution imaging. However, a high *NA*_*obj*_ is normally only possible with a short effective focal length and high system magnification. As such, high-resolution image information is captured only over a small field of view (FoV). The total image information, which can be quantified using the space-bandwidth product (SBP) measure [5], tends to decrease with increasing spatial resolution. Mechanical tiling can be used to extend the FoV. However, this approach is susceptible to blurring from stage drift and focus variations caused by specimen topology, as well as stitching artifacts arising from non-uniform illumination, exposure variations, and vignetting. Fourier ptychographic microscopy (FPM) is a computational imaging technique that overcomes this resolution-to-FoV trade-off by synthetically increasing the NA of the system (*NA*_*syn*_) using inclined spatially coherent illumination to shift the Fourier spectrum of the sample region captured by the microscope [6,7]. Iterative phase retrieval algorithms are used to recover a high-resolution complex image from a sequence of brightfield (BF) and darkfield (DF) images with overlapping spectra [8]. The resulting amplitude and phase images have a spatial resolution that depends on *NA*_*syn*_ = *NA*_*obj*_ + *NA*_*illum,max*_, where the maximum illumination NA is *NA*_*illum,max*_ = *n* sin *θ*_*max*_ and *θ*_*max*_ is the largest illumination angle [9]. FPM is a particularly promising approach for low-cost microscopy because high-resolution images can be captured using low-NA optics. By jointly estimating the sample and pupil function [10], the refractive imaging aberrations associated with low-cost optical components can be computationally corrected post-capture. Previous low-cost FPM systems used 3D-printed components [11], a smartphone sensor and illumination display [12], and a Raspberry Pi camera (and accompanying lens) [13]. However, these systems are limited in their flexibility, with light emitting diodes (LEDs) fixed in position, preventing axial adjustment for setting the passband overlap and minimizing vignetting. Additionally, the use of smartphones, optical elements, and cameras constrains the modularity and accessibility of these systems.

In this article, we report OpenFPM, a compact FPM platform that integrates a low-cost finite-conjugated objective lens and consumer-grade LED array into an open-source 3D-printed microscope body. Compared to previous implementations, OpenFPM provides flexible Fourier-space sampling via a translational LED array and offers improved modularity by supporting low-cost finite-conjugated objectives and widely available c-mount cameras. The OpenFPM microscope shown in Figure 1 was assembled from components fabricated with matte black polylactic acid (PLA) filament, using a Bambu Labs P1S fused deposition modelling (FDM) printer. The design of the microscope body was based on the OpenFlexure Project [1] version 7, which was modified for FPM. Designs for all 3D printed components, including the LED-array frame design, are available on GitHub [14]. Illumination, stage and objective positioning, and camera triggering were controlled using an open-source Python-based single-script graphical user interface, built on the Raspberry Pi 4 model B single-board microcomputer. Illumination was provided by a 16 × 16 LED array of individually addressable LEDs (WS2812B, BTF lighting). As indicated in Figure 1(a), the LED to sample height was adjustable in 10 mm increments using a pin-and-hole system. Prior to image acquisition, the LED array can be coarsely aligned manually, aided by realtime visualization of the overlapping coherent passbands in the Fourier spectrum of a suitably high-contrast sample, such as stained cells or a thin tissue section. Figure 1(b) shows an OpenFPM system, in which a commercial grade standard finite conjugate 10×/0.25 NA PLAN objective lens (Edmund Optics) with a working distance of 1.5 mm, and tube length of 160 mm forms an image of the specimen on the camera sensor. A plane aluminum steering mirror (Thorlabs) was used to reduce the overall height of the microscope. This system uses a monochrome CMOS camera (IDS UI-3060CP Rev.2, Sony IMX174LLJ-C), however the OpenFPM platform is compatible with any c-mount camera that supports hardware triggering for synchronization of illumination and image acquisition, using General-Purpose Input/Output pins on the Raspberry Pi. Each image was captured and stored on an external laptop for processing using this pipeline and the ThorCam software (64-Bit Windows version 3.7.0). Raw FPM images were acquired under sequential illumination from 177 LEDs, arranged in a circular pattern. The LED-sample plane distance was set to 80 mm to minimize the number of BF–DF transition images, thereby reducing the reconstruction artifacts produced by vignetting [15]. This enabled robust reconstruction of uniformly illuminated, high-resolution images with a 1 mm FoV, a nominal *NA*_*syn*_ of 0.92, and a 69% overlap between the Fourier space bands of images captured using adjacent LEDs. To quantify the spatial resolution of the FPM amplitude and phase images compared with conventional BF microscopy, an incoherent image was computed as the sum of all 177 raw images. With an exposure time of 200 ms per image, a complete FPM dataset consisting of 9 BF and 168 DF images was acquired in ∼40 s. For color FPM imaging, this acquisition was repeated for each RGB channel, with exposure times of 200 ms (red, 636 nm), 160 ms (green, 523 nm) and 100 ms (blue, 470 nm), using the maximum brightness for each LED, consistent with the camera sensor quantum efficiency. With a camera pixel size of 5.86 μm, the effective image pixel size for the 10× system is 0.586 μm, corresponding to a Nyquist spatial frequency limit of 853 cycles/mm, as shown in Figure 2(d). For conventional brightfield imaging, the incoherent cutoff frequency is given by 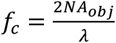, where *λ* is the illumination wavelength. As a result, the red images were oversampled, while the green and blue images were marginally undersampled. Chromatic focus shifts were corrected by repositioning the objective lens (using a Z-focus motor) between color channels. Based on the nominal values of *NA*_*obj*_ and *NA*_*syn*_ at 636 nm the depth of field (DoF) of the incoherent BF and reconstructed FPM amplitude and phase images was 12.52 μm and 1.39 μm respectively [16]. FPM images were reconstructed using the sequential quasi-Newton phase retrieval algorithm described by Tian et al. [17]. Reconstructions were performed using open-source software [18] in MATLAB (R2024a, MathWorks, Natick, MA) on a laptop equipped with an Intel Core i7 CPU (2.6 GHz, 32 GB RAM) and an NVIDIA RTX 2080 Super GPU (8 GB VRAM) running Windows 11. An angle self-calibration algorithm for LED correction described by Rogalski et al. [18] converged in approximately 2 min using GPU computation on a (200 pixel × 200 pixel) tile. To satisfy the spatial coherence constraints imposed by the van Cittert–Zernike theorem [19] and mitigate memory limitations [6], the raw image series (1936 pixels × 1216 pixels × 177 frames) was divided into 40 tiles (256 pixels × 256 pixels) with a 10% overlap. Six tiles were processed simultaneously using all the available CPU cores. Following reconstruction, tiles were stitched to form the full FoV, with complete reconstruction requiring 10-12 min per color channel. For RGB imaging, rigid registration [20] was applied to each tile to correct the lateral chromatic offsets between the channels prior to stitching. Final color images were generated in FIJI [21,22] using automatic brightness and contrast adjustment, with a manually defined background region of interest (ROI) used for white balancing. Image resolution was experimentally quantified under 636 nm illumination using a high-resolution Siemens star target (PAIR 1, Ready Optics, Calabasas CA, USA) [23]. Intensity measurements were extracted from 14 concentric rings on the Siemens star by applying a polar 360 degree transformation to the incoherent BF, FPM amplitude and FPM phase images in FIJI [21], seen in Figure 2(a-c). The Michelson contrast, 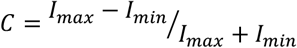, for each ring was determined by the average maximum (*I*_*max*_) and minimum (*I*_*min*_) intensity values for the spoke and space periods, as the spatial frequency increases towards the center of the star. Figure 2(d) shows these values plotted as a modulation transfer function (MTF). The resolution limit was quantified at 10% contrast corresponding to spatial frequencies, 526 cycles/mm (Incoherent BF), 1430 cycles/mm (FPM amplitude) and 1172 cycles/mm (FPM phase). Therefore, the full-pitch resolution limits were measured as 1.87 μm (Incoherent BF), 700 nm (FPM Amplitude) and 853 nm for FPM Phase, limited by phase reconstruction artifacts near the center of the star. Furthermore, the Abbe diffraction limit formula (*d* = ^*λ*^/*2NA*_*obj*_) was used to determine the effective conventional *NA*_*obj*_ ≈ 0.17. Additionally, the rewritten Abbe criterion for FPM, where the conventional condenser NA (*NA*_*cond*_) is replaced by *NA*_*illum,max*_, confirmed the computational *NA*_*syn*_ ≈ 0.74 and 0.90, for FPM phase and amplitude data respectively [24]. To demonstrate the different imaging modes possible with OpenFPM, the system was used to image a Giemsa-stained thin film blood smear [4]. Figure 3 shows representative (a) incoherent BF, (b) monochromatic FPM amplitude, (c) FPM phase, and (d) FPM color amplitude with (iv) a digitally magnified ROI without image registration. High-resolution, high-contrast FPM reconstructions enable visualization of the morphological features of red blood cells (RBCs) and white blood cells (WBCs) over an extended FoV [25]. The WBC shown in Figure 3(d) can be identified as a neutrophil due to its multi-lobed nuclear morphology. The FPM phase image (c) contains contrast information, with the WBC clearly distinguishable from surrounding RBCs owing to differences in the optical path length arising from variations in refractive index and cellular morphology. Important for clinical applications, including malaria diagnosis, the FPM RGB color amplitude image in (d) allows the distribution of the (dark purple) Giemsa-stain to be clearly visualized throughout the blood smear. The raw OpenFPM image sequence can be processed to render additional imaging modes including DF (Figure 3(e)) and Rheinberg contrast (Figure 3(f)) – a combination of BF and DF information – advantageous for edge detection and object visualization respectively. Finally, we investigated variations in image quality across the FoV by analyzing the pupil functions recovered for each reconstructed image patch. The retrieved pupil phase was decomposed into Zernike modes before the root mean square (RMS) wavefront error was calculated after removing the piston, x-tilt, and y-tilt terms. The heatmap in Figure 4(a) shows that the Strehl ratio (calculated from the RMS values using the Maréchal approximation [26]) varies between 0.82 and 0.33, with a low wavefront error and high image quality at the center of the image and a lower image quality and larger wavefront error in the lower left and upper right corners. Figure 4(b) shows the pupil phase associated with the lowest (green box) and highest (blue box) RMS wavefront error. Figure 4(c) shows the amplitudes of different modes ordered using the ANSI indexing scheme [27]. At the corner of the image, modes 5 (defocus) and 9 (coma) had the largest magnitudes. The pupil phase in the center is primarily comprised of circularly symmetric modes, including spherical aberrations.

**Fig. 1.**
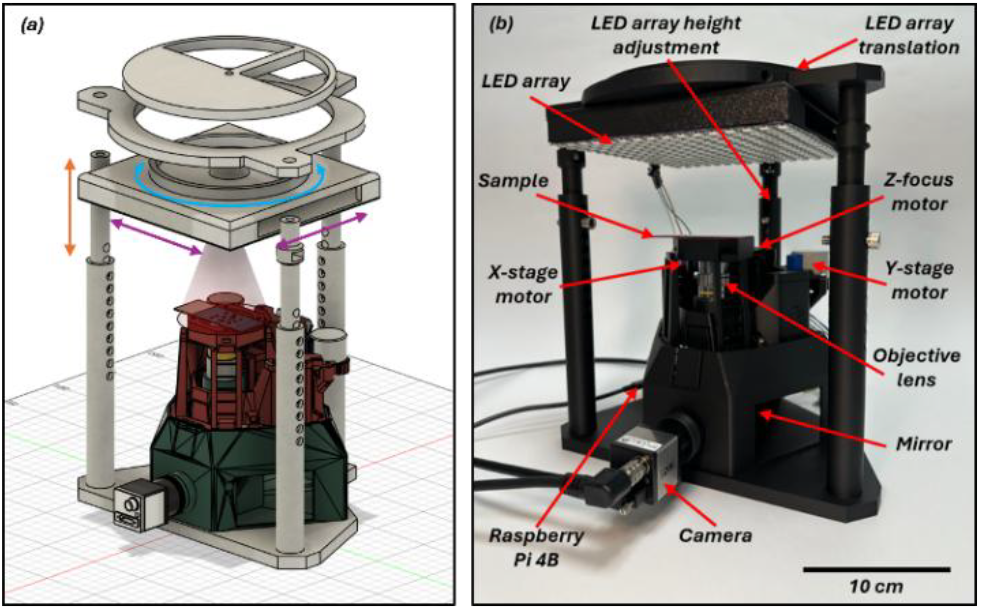
CAD model and 3D-printed implementation of the OpenFPM system. (**a**) Computer-aided design (CAD) model of the OpenFPM microscope, created in Autodesk Fusion 360 (version 2605.1.52), with arrows indicating the LED array degrees of freedom relative to the sample plane. Grey: original design, green: custom OpenFlexure, red: original OpenFlexure. (**b**) Photograph of the assembled 3D-printed microscope with key components labelled.

**Fig. 2.**
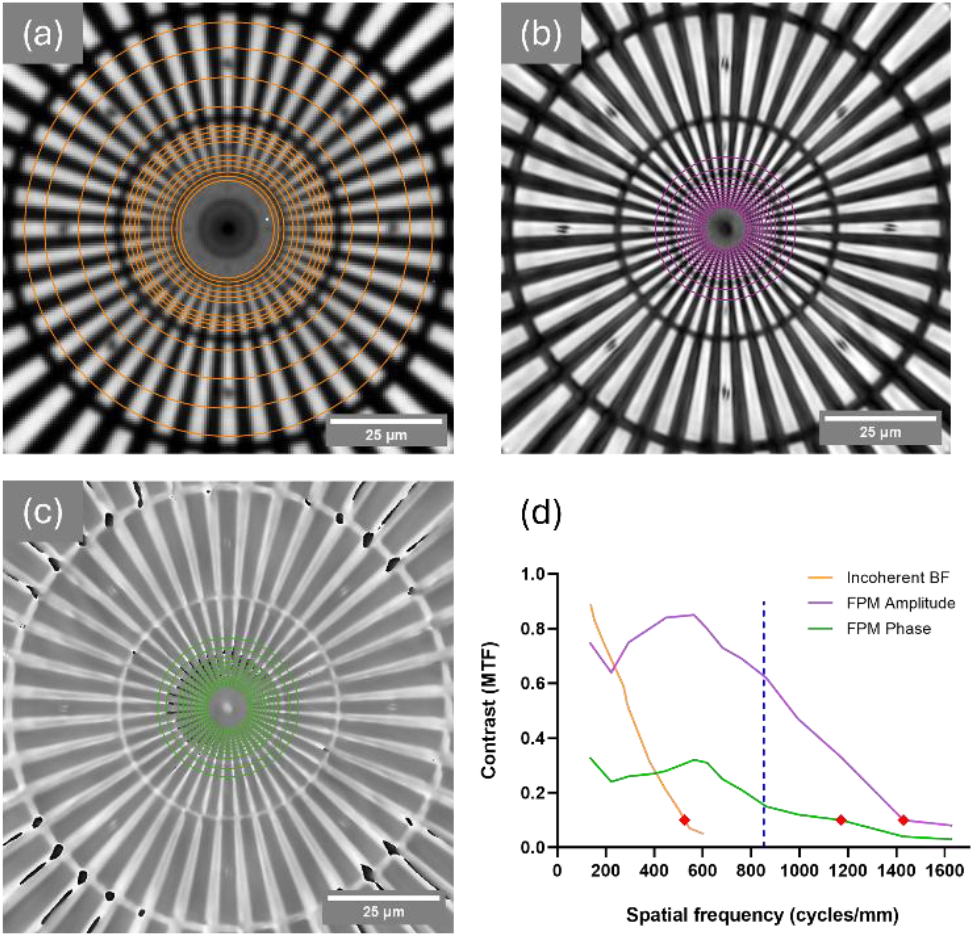
Experimental quantification of spatial resolution using Siemens star. (**a**)Incoherent BF ROI. (**b**) FPM Amplitude ROI. (**c**) FPM Phase ROI. (**d**) MTF of contrast levels and associated spatial frequencies measured from intensity data present in (**a-c**), Nyquist spatial frequency limit for 636 nm illumination (blue dotted line), and 10% contrast thresholds (red diamonds).

**Fig. 3.**
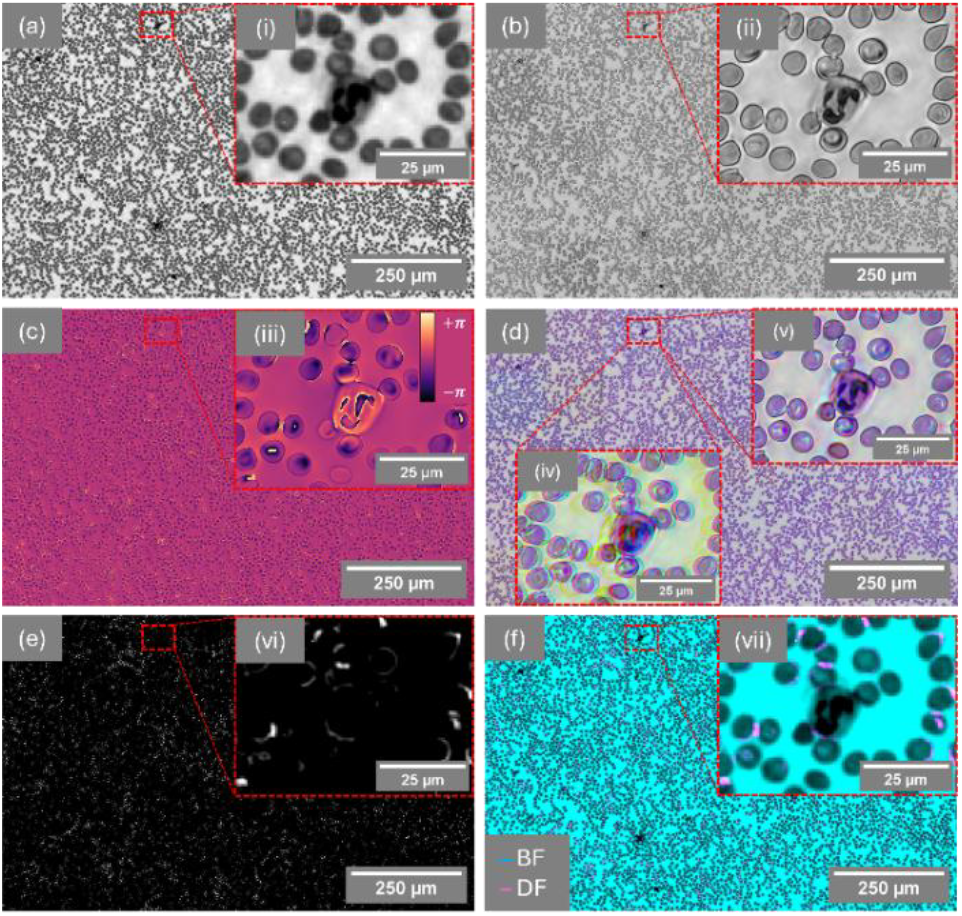
Imaging modalities using a Giemsa-stained blood smear. (**a**) Incoherent monochromatic BF. (**i**) Digitally magnified ROI of RBCs and WBC. (**b**)FPM amplitude reconstruction. (**ii**) ROI. (**c**) FPM wrapped phase contrast reconstruction (Lookup table: mpl-magma). (**iii**) ROI with colormap (radians). (**d**) FPM color amplitude reconstruction. (**iv**) ROI without image-registration and AWB (**v**) ROI with image-registration and AWB. (**e**) Monochromatic DF. (**vi**) ROI. (**f**) Rheinberg contrast [BF: cyan, DF: magenta]. (**vii**) ROI.

**Fig. 4.**
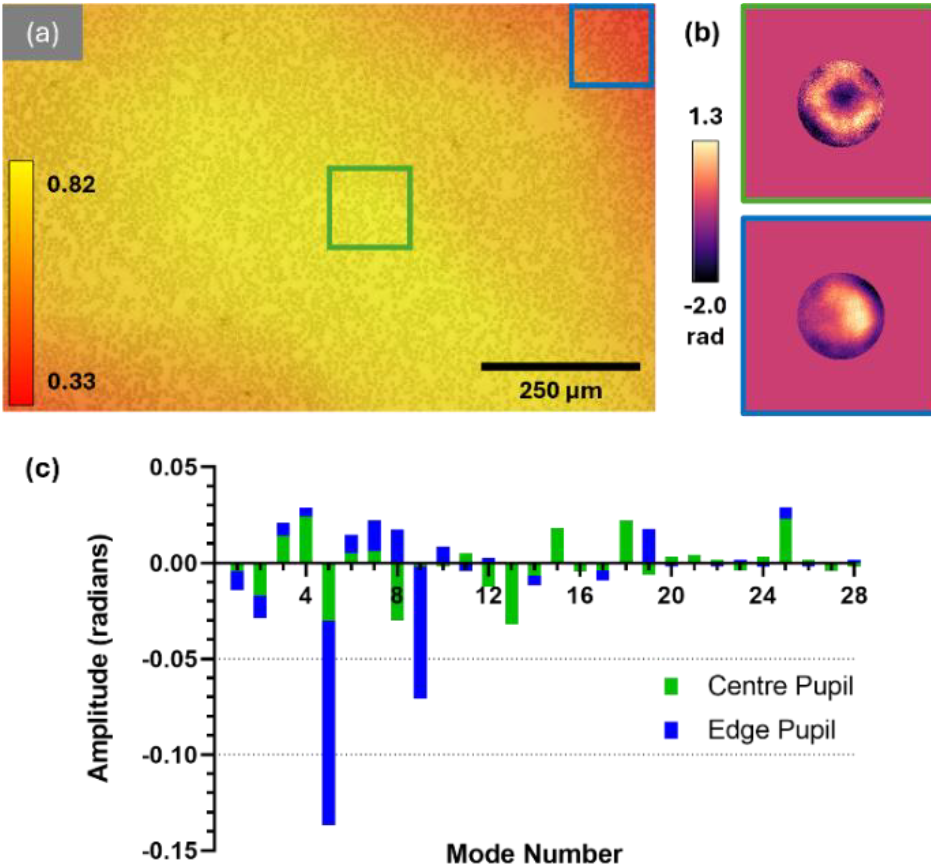
Recovered phase of pupil functions, Zernike decomposition and Strehl ratio heatmap over the full FoV. (**a**) Two dimensional Strehl value heatmap over the full FoV FPM amplitude image. (**b**) Two ROIs of the phase pupil functions with the highest (green) and lowest (blue) Strehl value. (**c**) Zernike decomposition of the two pupil function error maps. Modes 5 (defocus) and 9 (coma) have the large magnitudes at the edge of the FoV.

The architecture of OpenFPM lowers the barrier to LED-array based modular hardware, lowering costs, and enabling flexibility of optics, illumination geometry, and software. This approach allows high-resolution, large-FoV imaging, digital aberration correction, and multimodal imaging to be achieved within a compact system, supporting the development of computational microscopy techniques for developers and end users. Experimental simplicity and accessibility make OpenFPM attractive for many biomedical applications, including adoption in resource-constrained settings. Within this system, the illumination source remains a key element of imaging performance in FPM. Whilst the current OpenFPM design uses a planar LED array, studies have demonstrated that quasi-dome [28] and hemispherical [29] geometries offer advantages, including an increased signal-to-noise ratio in darkfield images, reduced image acquisition times and higher effective spatial resolution. Therefore, the integration of new low-cost LED illuminators is a particularly promising avenue for further development of the OpenFPM platform.

## Funding

LDW and LC were supported by the Engineering and Physical Sciences Research Council (EP/Y528833/1) and UK National Measurement System. LMR and GM were supported by Leverhulme Trust. LMR was funded by the University of Glasgow. CB was supported by an award from the UCL CS Innovation Seed Fund. MS acknowledges funding from the Life Sciences and Health Program of the UK National Measurement System and the NIHR UCLH Biomedical Research Centre (award 187809). GM was supported in part by the Biotechnology and Biological Sciences Research Council (BB/V019643/1, BB/X005178/1, BB/T011602/1, and BB/W019032/1).

## Disclosures

The authors declare no conflict of interest.

## Data Availability

Data underlying the results presented in this paper are available in Ref. [14].

## Notes

https://github.com/YoItsLewis/Open-Fourier-Ptychographic-Microscopy

### Competing Interest Statement

The authors have declared no competing interest.

## REFERENCES

1. J. T. Collins, J. Knapper, J. Stirling, et al., Biomed. Opt. Express 11, 2447 (2020).

2. L. M. Rooney, J. Christopher, B. Watson, et al., Adv Materials Technologies 9, 2400043 (2024).

3. M. Del Rosario, H. S. Heil, A. Mendes, et al., Advanced Biology 6, 2100994 (2022).

4. J. Christopher, R. Craig, R. E. McHugh, et al., Journal of Microscopy 298, 274 (2025).

5. A. W. Lohmann, R. G. Dorsch, D. Mendlovic, et al., J. Opt. Soc. Am. A 13, 470 (1996).

6. G. Zheng, R. Horstmeyer, and C. Yang, Nature Photon 7, 739 (2013).

7. J. W. Goodman, (Roberts and Company Publishers, 2005).

8. L.-H. Yeh, J. Dong, J. Zhong, et al., Opt. Express 23, 33214 (2015).

9. X. Ou, R. Horstmeyer, G. Zheng, et al., Opt. Express 23, 3472 (2015).

10. X. Ou, G. Zheng, and C. Yang, Opt. Express 22, 4960 (2014).

11. S. Dong, K. Guo, P. Nanda, et al., Biomed. Opt. Express 5, 3305 (2014).

12. K. C. Lee, K. Lee, J. Jung, et al., ACS Photonics 8, 1307 (2021).

13. T. Aidukas, R. E T. Aidukas, R. Eckert, A. R. Harvey, et al., Sci Rep 9, 7457 (2019).

14. L. D. Walker, “Open Fourier Ptychographic Microscopy,” GitHub (2026), https://github.com/YoItsLewis/Open-Fourier-Ptychographic-Microscopy.

15. A. Pan, Chao Zuo, Yuege Xie, et al., Optics and Lasers in Engineering 120, 40 (2019).

16. R. Claveau, P. Manescu, M. Elmi, et al., Biomed. Opt. Express 11, 215 (2020).

17. L. Tian, X. Li, K. Ramchandran, et al., Biomed. Opt. Express 5, 2376 (2014).

18. M. Rogalski, P. Zdańkowski, and M. Trusiak, Bioinformatics 37, 3695 (2021).

19. R. Eckert, Z. F. Phillips, and L. Waller, Appl. Opt. 57, 5434 (2018).

20. P. Thevenaz, U. E. Ruttimann, and M. Unser, IEEE Trans. on Image Process. 7, 27 (1998).

21. J. Schindelin, I. Arganda-Carreras, E. Frise, et al., Nat Methods 9, 676 (2012).

22. S. Preibisch, S. Saalfeld, and P. Tomancak, Bioinformatics 25, 1463 (2009).

23. R. Horstmeyer, R. Heintzmann, G. Popescu, et al., Nature Photon 10, 68 (2016).

24. F. Xu, Z. Wu, C. Tan, et al., Cells 13, 324 (2024).

25. M. Shaw, R. Claveau, P. Manescu, et al., The Journal of Pathology 255, 62 (2021).

26. V. N. Mahajan, J. Opt. Soc. Am. 73, 860 (1983).

27. Paul Fricker, “Zernike polynomials,” MATLAB Central File Exchange (2026), https://uk.mathworks.com/matlabcentral/fileexchange/7687-zernike-polynomials.

28. Z. F. Phillips, R. Eckert, and L. Waller, in Imaging and Applied Optics 2017 (3D, AIO, COSI, IS, MATH, pcAOP) (OSA, 2017), p. IW4E.5.

29. M. G. Mayani, N. Hussain, D. W. Breiby, et al., Opt. Eng. 64, (2025).

